# Structural stability determines evolutionary stability in mutualistic model ecosystems

**DOI:** 10.1101/2024.09.04.611292

**Authors:** Miguel Lurgi, Alberto Pascual-García

## Abstract

Understanding the factors that influence the persistence and stability of complex ecological networks is a central focus of ecological research. Recent research into these factors has predominantly attempted to unveil the ecological processes and structural constraints that influence network stability. Comparatively little attention has been given to the consequences of evolutionary events, despite the fact that the interplay between ecology and evolution has been recognised as fundamental to understand the formation of ecological communities and predict their reaction to change. In light of current environmental challenges, there is a compelling need for a quantitative framework to predict biodiversity loss under environmental perturbations while accounting for evolutionary processes.

We extend existing mutualistic population dynamical models by incorporating evolutionary adaptation events to address this critical gap. We relate ecological aspects of mutualistic community stability to the stability of persistent evolutionary pathways. Our findings highlight the significance of the structural stability of ecological systems in predicting biodiversity loss under both evolutionary and environmental changes, particularly in relation to species-level selection. Notably, our simulations reveal that the evolution of mutualistic networks tends to increase a network-dependent parameter termed critical competition, which places systems in a regime in which mutualistic interactions enhance structural stability and, consequently, biodiversity.

This research emphasizes the pivotal role of natural selection in shaping ecological networks, steering them towards reduced effective competition below a critical threshold where mutualistic interactions foster stability. The outcomes of our study contribute to the development of a predictive framework for eco-evolutionary dynamics, offering insights into the interplay between ecological and evolutionary processes in the face of environmental change.

## Introduction

Unveiling the drivers of persistence and stability of complex ecological networks remains a primary focus of ecological research. Seminal work by MacArthur in the 1960s suggested a positive relation between the complexity of ecological interactions and stability [1]. A decade later, May challenged this notion with mathematical proof of an opposing result, showing the existence of a limit in the size and connectivity of an ecological system for it to be stable [2]. May’s work also suggested that mutualistic interactions are less compatible with the stability of large ecosystems compared with other interaction types: ‘It is tempting to speculate that such community stability considerations may play a role in explaining why mutualism *is a fascinating biological topic but its importance in populations in general is small’* (May in [3] quoting Williamson). This claim seems paradoxical considering the significant biomass hosted by flowering plants across the world’s biomes and the extensive biodiversity of their pollinators [4]. These systems are fundamentally supported by mutualistic interactions. Furthermore, mutualistic interactions in macroscopic [5], as well as in microscopic organisms [6] are widespread across many ecological contexts.

Lotka-Volterra models of interacting populations dynamics have played a pivotal role in our theoretical understanding of community persistence. A central focus of research grounded on these models has been on comparing the stability of purely competitive communities with those incorporating mutualistic interactions. Reported negative effects of mutualisms on community persistence have been linked to the absence of saturating responses for mutualistic effects. Ironically, as early as in 1979 Goh showed that non-linear models of mutualism were more tractable than competitive or prey-predator ones [7]. Nowadays, it seems clear that non-linear responses lead to stable systems sustaining biodiversity [8, 9], a result that is robust even in the face of invasions by alien species [10].

Another critical factor influencing the relationship between mutualisms and community stability is the consideration that, although models with random parameters are extremely valuable as null models (see e.g. [11]), explanatory models often require non-random parameters. Indeed, interest in the study of mutualistic interaction networks surged partly from field experiments reporting non-random nested patterns in the organization of their links [12]. These observations prompted the development of theoretical approaches investigating the importance of network topology for the stability of simulated mutualistic communities.

While some researchers have reported a positive relationship between nestedness and the stability of mutualistic communities [9, 13], others have suggested a more prominent role of other topological properties such as the connectance or the degree distribution within these networks [14]. These discrepancies have led to the suggestion that fair comparisons of the stabilizing effects of different topological properties should only be made after ensuring the feasibility of the systems being compared [15, 16, 17]. This is particularly important for comparisons between systems showing substantial differences in the number of parameters, e.g. purely competitive communities vs those considering mutualistic and competitive interactions together. Random parameter value selection confounds feasibility and stability, a distinction not always acknowledged [18], making systems with fewer parameters appear more stable.

This focus on structural stability, specifically the stability of feasible and locally stable systems against perturbations on system parameters [15, 17, 9], led one of us & Bastolla to find that mutualistic interactions favor biodiversity only under certain conditions. The strength of interspecific competitive interactions among species within the same trophic level (e.g. plants or pollinators) must be smaller than a threshold termed critical competition for a mutualistic system to be more structurally stable than a purely competitive system [16]. The value of topological properties making these systems more structurally stable also changes depending on whether the strength of interspecific competition is above or below this critical value. Crucially, network connectivity increases stability when competition is below this critical value but has the opposite effect when competition is above it. Nestedness on the other hand has a notable positive effect on stability either when competition surpasses the critical value, or below the critical value only if the connectance is low. It has been claimed that these findings explained previous discrepancies regarding the effect of mutualisms on complex ecosystems and the topological properties relevant for biodiversity maintenance [19, 20].

Recent work investigating the evolutionary assembly of mutualistic networks argue for the existence of mutualistic regimes favoring biodiversity. These efforts have revealed the existence of structural patterns such as nestedness [21] or modularity [22] which, as it was the case for dynamical stability, may also depend on the assumptions made. For example, the absence of explicit ecological dynamics [23], a focus on indirect mutualism [24], or the explicit consideration of demographic stochasticity [25] or species traits [26] have led to different conclusions on the emergence of structural properties in mutualistic networks. As highlighted above for the case of structural stability, another important consideration when comparing the outcome of different studies is the choice of parameters, which may influence the expected ecological behaviour [10]. These potential sources of discrepancy across studies highlight the need for a unified eco-evolutionary framework considering the minimal set of ingredients capable of generating expected structural outcomes of mutualistic networks depending on different choices of mechanisms and parameters. Although we are still far from achieving this goal, a step forward is to explore whether it is possible to predict the evolutionary outcome (expected biodiversity) from the properties of a starting unperturbed system. In particular, how different considerations regarding the connectivity and endurance to change of these systems may influence such predictions. There is increasing interest in relating environmental conditions and network properties (see e.g.[27, 28]), given the rapidly changing environmental conditions we are witnessing.

In this work, using a theoretical eco-evolutionary framework, we explored the effects that the combined action of evolutionary events (in the form of changes in species interactions and subsequent species-level selection) and environmental fluctuations have on the biodiversity of mutualistic systems (Fig. 1a). More specifically, we asked: i) whether it is possible to predict biodiversity loss without species reintroduction; ii) to what extent the behavior observed for the structural stability of the mutualistic network, i.e., mutualistic benefits depending on the value of intraguild competition relative to a critical threshold, still holds when evolutionary events are considered (illustrated in Fig. 1b); and, as a corollary, iii) what mutualistic regime (beneficial or detrimental for stability) systems tend to evolve towards (interspecific competition below or above critical competition).

**Figure 1:**
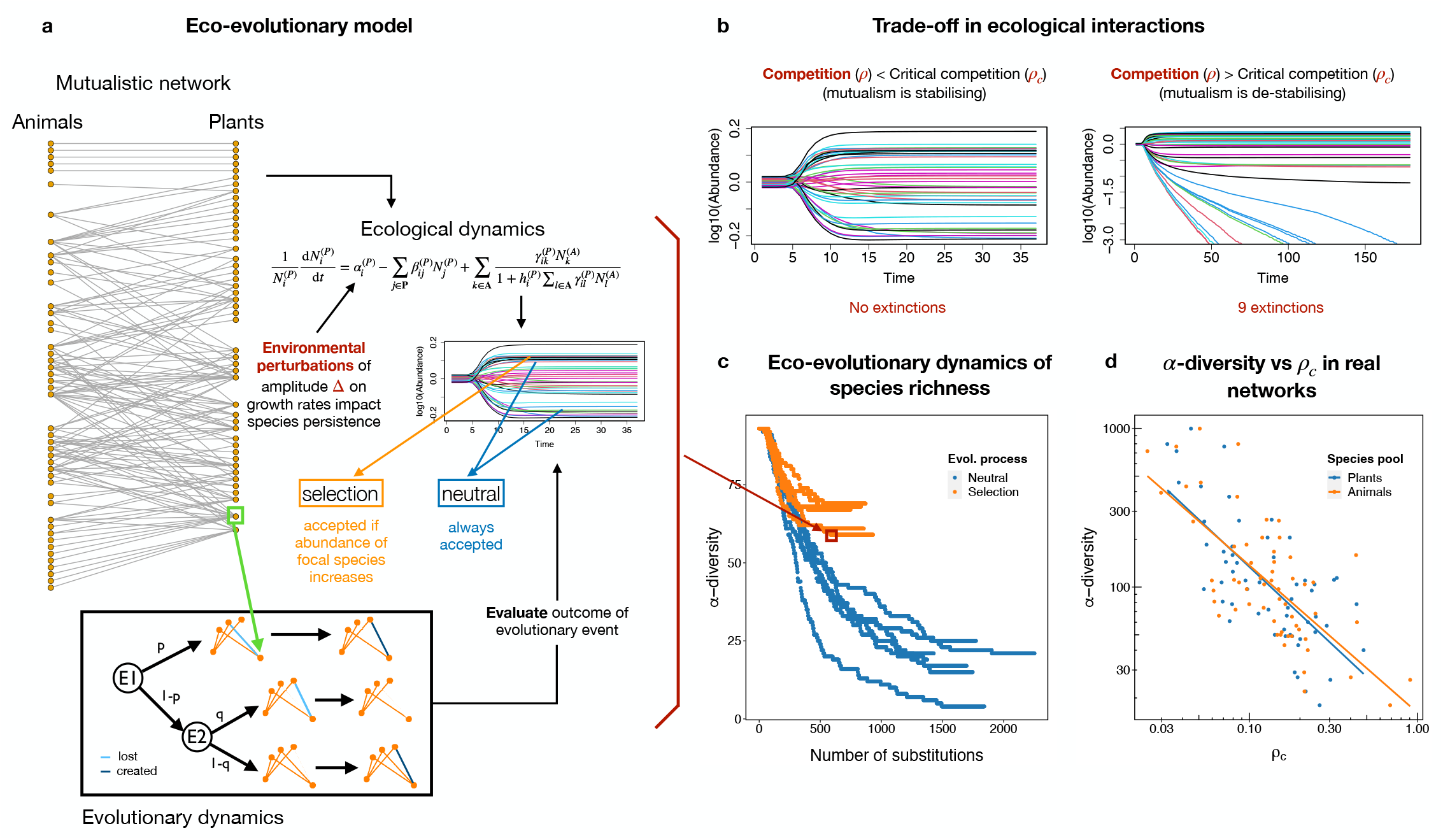
Eco-evolutionary model and critical competition *ρ*_*c*_. (a) Schematic representation of the model. At each step of the simulation, a randomly-selected species (green square) is subjected to an evolutionary event (box “evolutionary dynamics”). Evolutionary events are of two types: link swap (E1) or creation / loss (E2). The probabilities of events (*p, q* respectively) are indicated. The effect of the event on the network is evaluated by running the ecological dynamics, where environmental perturbations may be added. Acceptance of the event depends on whether selection is considered. If selection is considered, the change is accepted only if it enhances the fitness of the modified species (i.e. there is an increase in its biomass). In the case of “neutral” selection all changes are accepted, regardless of their effect on species or the community. (b) Simulated time series of species abundances illustrating the trade-off between the interspecific competition parameter *ρ* and the critical competition *ρ*_c_. (c) Example of eco-evolutionary dynamics showing six trajectories for the neutral (blue) and selection (orange) scenarios. The evolutionary equilibrium is achieved when no extinctions are observed throughout 350 substitutions (i.e. accepted evolutionary events). (d) log-log representation of the critical competition against the total biodiversity for a set of real plant-pollinator networks extracted from the Web of Life. The power-law relationships for both plants and animals have an exponent ∼ − 0.5. By means of the trade-off illustrated in (b), observing positive mutualistic effects in large communities demand decreasing the intraguild interspecific competition to favour coexistence.

Our simulations demonstrate that less competitive systems are not only more structurally stable but also more robust against changes brought about by evolutionary events. Crucially, estimation of the structural stability of the system before the evolutionary process is started is a good predictor of the evolutionary outcome in the presence of selection, especially when nuanced environmental perturbations are considered. We show that the main mechanism sustaining biodiversity throughout evolutionary time is an increase in the system’s critical competition. Consequently, the system increases its chances of stability in the face of environmental perturbations while maintaining mutualistic interactions in the regime where mutualism positively contributes to structural stability, promoting biodiversity maintenance. We would thus expect ecological networks shaped by natural selection to exhibit a tendency to keep effective competition below the critical value of competition to maintain biodiversity and withstand perturbations.

## Results

### The interplay between effective and critical competition determines stability and diversity

We developed an eco-evolutionary model for mutualistic networks by coupling an ecological community dynamics model with an evolutionary model of species interactions, illustrated in Fig. 1a. The model proceeds as follows: we begin by setting the system at a feasible and dynamically stable state. Next, we introduce an evolutionary event by randomly selecting a species that will either: i) swap one of its interactions with another randomly selected species with probability *p*; ii) establish a new interaction with probability (1 *− p*)*q*; or iii) remove it with probability (1 *− p*)(1 *− q*). Unless otherwise is stated, we present simulations for *p* = *q* = 0.5 (see Methods and Suppl. Materials for other scenarios). Once the change is introduced, small perturbations emulating environmental noise might be introduced as uniform changes of amplitude Δ on the growth rates *α*_*i*_ of all species in the system. The resulting model is then integrated until equilibrium. Once the new equilibrium is achieved, the process is iterated by introducing a new evolutionary event, environmental perturbation, and integrating. In addition, we may consider the presence of selection if the species selected to introduce en evolutionary change must increase in biomass for the change to be accepted (the mutation becomes a substitution). If selection is absent we say the scenario is “neutral”, meaning that all mutations become substitutions independently of the fate of the species targeted. Note the abuse of language in the term *neutral*, since we do not intend to imply that the mutation does not have an effect on fitness, but that it is fixed independently of its effect.

The ecological model considered that mutualistic interactions between two pools of species (e.g. plants indexed *i* and their pollinator animal species indexed *k*) were encoded in a network of interactions *γ*_*ik*_ with a mean interaction strength *γ*_0_ (see Methods for a full description and Fig. 1 for a schematic representation). Competitive interactions within each pool were encoded in mean-field matrices *β*_*ij*_ defined with a parameter *β*_0_ for intra-specific and *ρ* for inter-specific competition (*β*_0_ > *ρ*) (see Fig. 1a and Methods). It is possible to rewrite the system containing both competitive and mutualistic interactions (and/or predatory, see [29]) as a purely competitive system, in which non-competitive interactions are effectively incorporated as variations in the *bare* interspecific competition *ρ*, becoming an *effective competition* parameter *ρ*_eff_. The most important advantage of the simplification provided by the formalism derives from its ecological interpretability, partly thanks to the relationship between the stability of the effective system and the whole system [30]. For instance, for mutualistic systems, it has been verified that *ρ*_eff_ < *ρ*, meaning that mutualistic interactions effectively reduce the interspecific competition [9]. The effect that specific network topologies have on the stability of the system can also be explored similarly [16]. Crucially, mutualistic benefits on structural stability were observed only if the effective interspecific competition parameter *ρ*_eff_ was lower than a threshold, *ρ*_c_, termed *critical competition* (Fig. 1b) [16].

In this work, and consistent with previous results, we observed that the relationship between *ρ*_eff_ and *ρ*_c_ determined different scenarios in which evolutionary changes showed a distinct behaviour depending on whether selection was considered or not. In Fig. 1c, we illustrate this for different runs of a simulation in which the same starting network was evolved under both neutral and selection scenarios. Evolution under selection consistently converged earlier to systems with a higher diversity. Since in real networks a negative relationship between diversity and critical competition has been observed (Fig. 1d), an open challenge is to derive the expected relationship between their effective and critical competitions to theoretically understand the mechanisms behind their stability. We aim to to tackle this challenge by investigating the effect that evolutionary events and environmental perturbations have on *ρ*_eff_ and *ρ*_c_, which influence the structural stability and, in turn, the diversity of the system.

### Structural stability predicts diversity at evolutionary equilibrium

We started by selecting a real network from the dataset available from the Web of Life (www.web-of-life.es/) to be evolved. We chose a network with approximately the same number of plant (47) and animal species (46) and connectance close to the median of the entire set *κ* = 0.069. By using the same initial mutualistic network across model simulations we could fairly compare the effect that the different ecological and evolutionary parameters had on the evolutionary outcome. We present in Suppl. Fig. 1 results for the evolution of randomly-generated networks, finding similar results.

We ran evolutionary simulations varying the value of intraguild interspecific competition *ρ* and the level of environmental perturbations from Δ = 0 (no perturbations: any speciess loss should be attributed to evolutionary events alone) to Δ = 0.1 (see Methods for details). The maximum value of Δ (0.1) was chosen because it determines the structural stability of the most vulnerable system (the one with highest competition, *ρ* = 0.15). The structural stability was defined as the minimum amplitude Δ of the perturbation on species’ growth rates leading to extinctions. We termed this value of the amplitude the critical perturbation Δ = Δ_c_ which can be estimated analytically, and verified numerically with simulations (see Methods and Ref. [16] for details). Unless otherwise is stated, for computational convenience we will use the analytical calculation. Therefore, Δ = 0.1 was close to the Δ_c_ of the less structurally stable system.

We found that the lower the interspecific competition, the higher the evolutionary stability, measured as species persistence (Fig. 2a). Interestingly, the specific relationship between diversity and competition changes depending on the presence and magnitude of environmental perturbations and on the existence of selection. In the absence of perturbations, the dependence of the final diversity of the system on the interspecific competition *ρ* is weaker than if perturbations are present (Fig. 2a). This dependence is, however, more nuanced under selection, with interspecific competition having little effect on biodiversity in neutral simulations (i.e. no selection) with strong environmental perturbations. The relative values between selection and neutral simulations also change depending on the specific value of the environmental perturbations, Δ. Selection is capable of preserving more diversity than neutral random change except for scenarios of relatively high interspecific competition in the absence of perturbations (Δ = 0). For high levels of perturbation, the loss of species observed for neutral simulations was up to ∼ 5-fold higher than in the presence of selection (forΔ = 0.1 and *ρ* = 0.0125).

**Figure 2:**
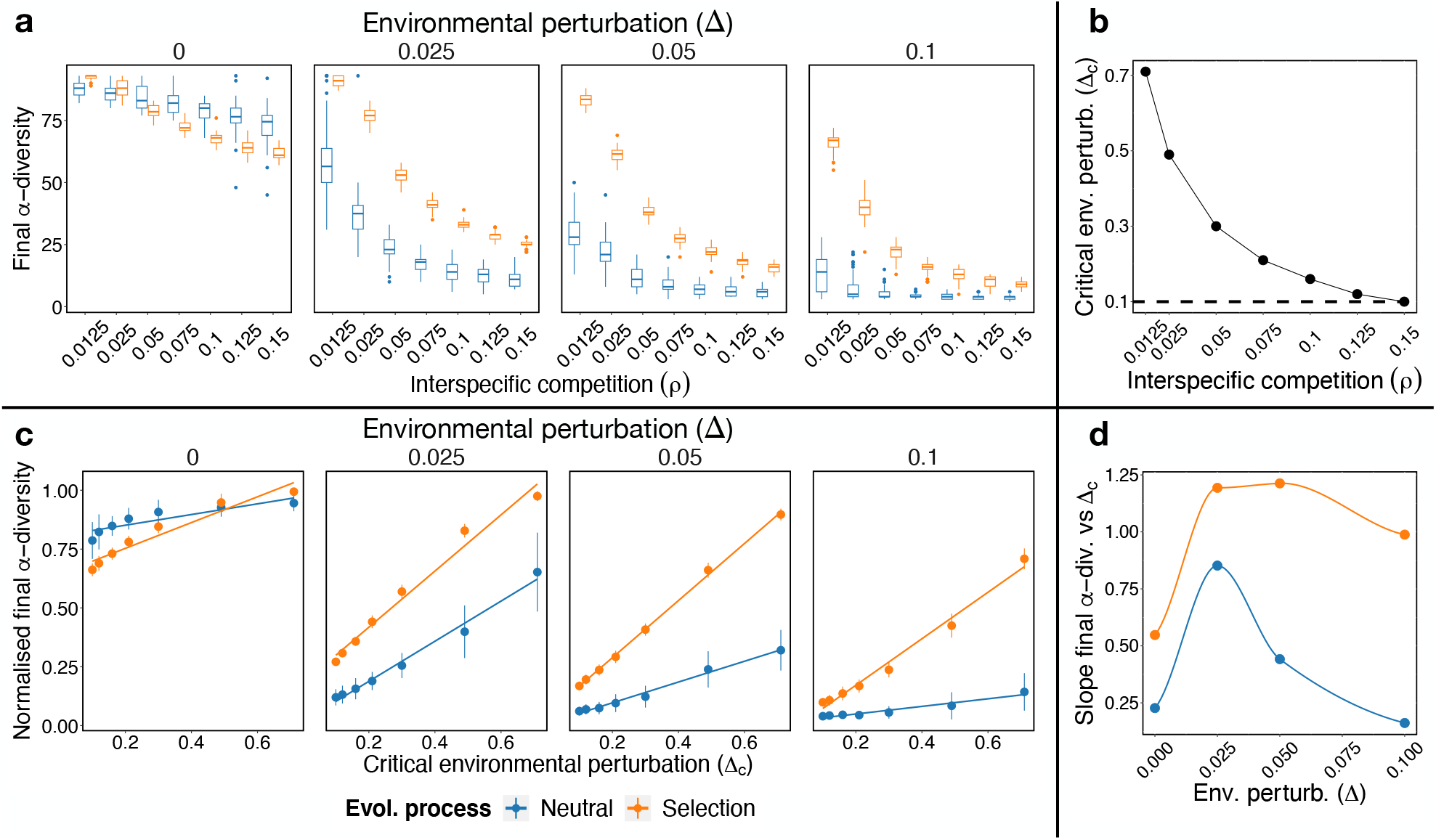
Structural stability predicts diversity at the evolutionary equilibrium. a) Biodiversity of the eco-evolutionary process at equilibrium for different values of the interspecific competition *ρ* and environmental perturbation Δ. Each panel represent the outcome of the evolutionary process for different values of Δ. In boxplots, box limits represent interquartile range (IQR) and the horizontal line the median value of the final species richness (*α*-diversity) across 50 simulations. Vertical lines (i.e. whiskers) show the +/- 1.5 IQR values. Points represent the outliers. b) Predicted structural stability (critical environmental perturbation Δ_c_) for the starting network for each value of interspecific competition *ρ*. c) Relationship between structural stability and final biodiversity, normalized by the starting diversity before the evolutionary process. Slope of the relationship shown in c) for different values of environmental perturbation. The closer the slope is to one the more accurate the prediction. Ordinates are not significantly different from zero for Δ = 0.1.

In Suppl. Materials, we extended the values of Δ explored by setting perturbations relative to the Δ_c_ of each system (i.e. Δ = *δ*Δ_c_ with *δ* ∈ [0, 1)). We found that more competitive systems still lose more species for the same relative perturbation (i.e. same value of *δ*), even though the absolute perturbation Δ is much higher for less competitive systems (Suppl. Fig. 2a). Moreover, we found that for each interspecific competition *ρ* value there is a characteristic environmental perturbation 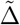 in which the relative difference in final *α*−diversity between simulations under selection and neutral scenarios peaked, with both quantities related as 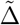 ∼ exp(−ρ) (Suppl. Fig. 2b-d). Therefore, given *ρ* it is possible to estimate the region of environmental perturbations amplitudes in which selection effects are stronger.

To quantify the agreement between evolutionary and structural stability, we compared the *α* diversity persistence (our proxy of evolutionary stability) against the predicted structural stability, Δ_c_, shown in Fig. 2c (see Methods). Notably, the evolutionary stability go hand in hand with structural stability showing a similar trend, with less competitive networks (i.e. those with higher Δ_c_; Fig. 2b) losing less species than those with high competition.

Moreover, representing the final diversity against the structural stability (Fig. 2c) we observed that structural stability had a good qualitative agreement in all scenarios, following significant linear trends in all cases. Moreover, as soon as environmental perturbations were present (Δ > 0), structural stability was a good predictor of the final diversity, with slopes approaching one (note that final diversity was normalized by the starting diversity to get a more direct interpretation of the slopes, shown in Fig. 2c). In Fig. 2d we summarise variations in the slopes of these relationships across the gradient of environmental perturbation

Δ. In the absence of environmental perturbation, structural stability underestimates the final expected diversity independently of the presence of selection, especially for highly competitive systems (lower structural stability; Fig. 2b). This is illustrated by the slope of the relationship in 2c being lower than one (Fig. 2d). For intermediate levels of perturbation, structural stability tends to overestimate the final diversity in the scenario under selection, and to underestimate in the neutral case. For high levels of perturbation, on the other hand, it tends to accurately predict the final diversity (slope tends to one) in the presence of selection, and strongly underestimate it for neutral simulations. In summary, structural stability qualitatively describes the evolutionary outcome, and it is a strong predictor of biodiversity in simulations under selection, in particular in the presence of higher levels of environmental perturbations.

It is important to note that structural stability predictions consider how the environmental perturbations acting on the growth rates of the starting system affects the persistence of the species in the system. On the other hand, evolutionary events affect the interactions between species, either swapping a randomly selected interaction between two species, introducing a new one, or removing a randomly selected interaction. Moreover, structural stability is defined on the starting system but, during the evolutionary process, the system will lose species and, hence, the structural stability of the evolved system changes. Therefore, the starting structural stability of the system sets the boundaries within which evolutionary biodiversity loss will lie.

### Biodiversity decline follows increasing biomass and connectance

To gather deeper insights into the impacts of evolutionary change on community and network structure, we selected specific simulations for further analysis. We focus on the two extreme scenarios of environmental perturbations (Δ = {0, 0.1}) and of interspecific competition (*ρ* = {0.0125, 0.15}), and assess their evolutionary trajectories in detail.

In the absence of environmental perturbation (Δ = 0) the qualitative behaviour for the loss of biodiversity was different for low and high competition (Fig. 3). For low competition under selection the system stabilized very quickly, conserving most species, while neutral simulations took longer times to stabilize, losing more species. On the contrary, for high competition the system under selection lost species faster but stabilised after ∼ 1500 substitutions, while the neutral simulations had a slower decay, and they were not stabilised after the limit of 2000 substitutions. For high environmental perturbations (Δ = 0.1), the loss of species was exacerbated for both low and high competition, with both neutral and under selection simulations having a very similar behaviour for high competition (Fig. 3). This is due to the fact that, for this parameterization, the environmental perturbation corresponds to the critical threshold of environmental perturbation, Δ_c_, hence being highly disruptive.

**Figure 3:**
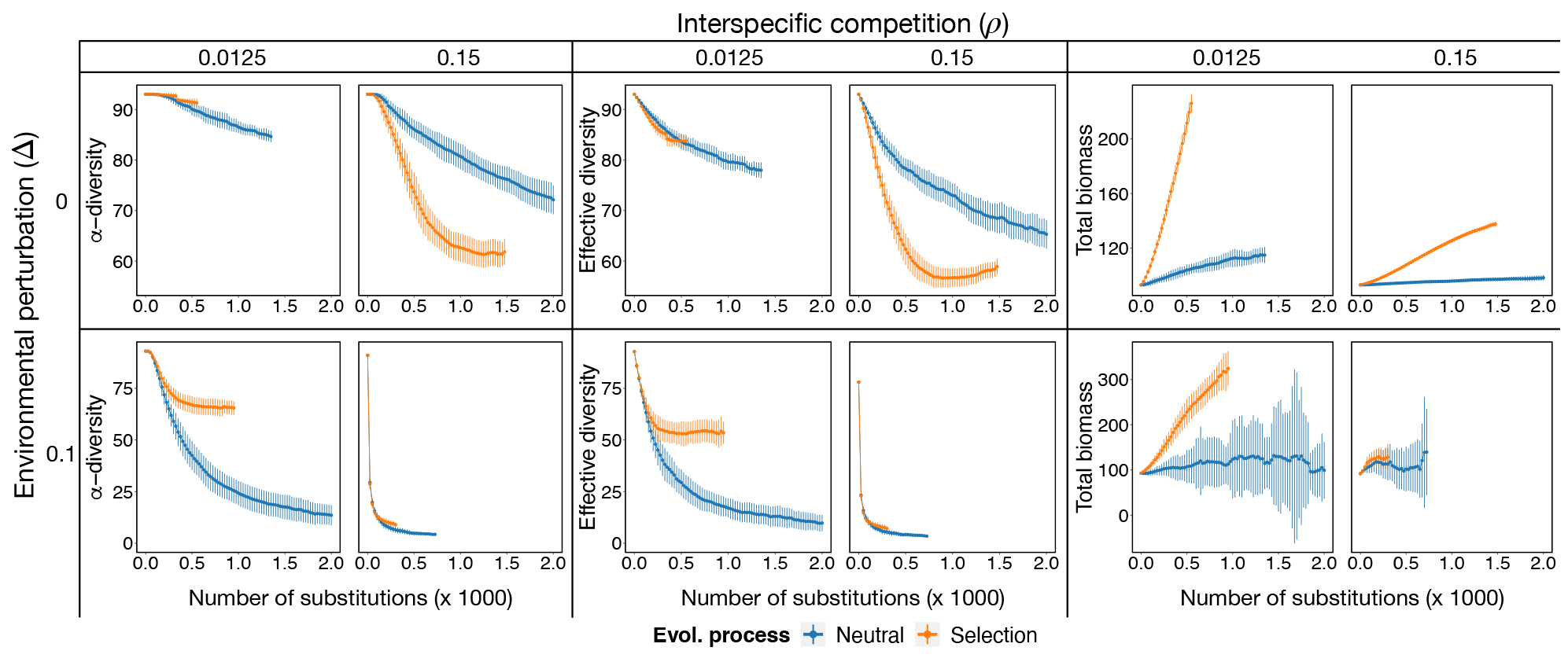
Disruptive evolution prompts biodiversity loss and biomass increase. Mean *α*-diversity (i.e. species richness) (left), effective diversity (middle) and biomass (right) throughout eco-evolutionary trajectories (halted after stability in number of species is achieved over 350 evolutionary steps) for different values of environmental perturbation Δ (rows) and interspecific competition *ρ* (columns). Mean values (points) and standard deviation (vertical lines) are shown every 25 evolutionary steps (i.e. fixed mutation or substitutions) across 50 independent simulations for selection (orange) and neutral (blue) scenarios.

In the presence of selection, the loss of biodiversity went hand in hand with a strong increase in total community biomass, except for scenarios of high environmental perturbations and high competition. An increase was expected, as the condition for selection is that the species subjected to evolutionary change must increase in biomass for the change to be accepted. Therefore, without perturbations, a faster loss of biodiversity under selection may be due to a destabilising effect of mutualistic interactions when biomass increases, as shown below. Notably, biomass was distributed unevenly across species in all systems, with some species concentrating more biomass, as indicated by a lower effective diversity compared to species *α* diversity (Fig. 3).

Biodiversity loss and biomass increase were accompanied by a significant increase in the connectance of the mutualistic network (see Fig. 3 and Suppl. Fig. 4). In the absence of environmental perturbations the increase was clearly a selected feature, as evidenced by the well-separated trend with respect to neutral simulations. When high environmental perturbations were present however, the increase in connectance seemed to be a collateral effect arising from the loss of species under high interspecific competition, with weak evidence for selection when competition was low. To confirm this interpretation, we ran simulations for two additional values of the parameter *p*, which controls the probability of performing an interaction insertion or deletion vs. a swap in the evolutionary events. When the probability of inserting new links was low and interactions rewiring (i.e. swaps) was favoured (*p* = 0.9), there was a more dramatic loss of biodiversity under selection (Suppl. Fig. 3a). Therefore, under this scenario, the increase in connectance was mainly driven by species loss (Suppl. Fig. 3). On the contrary, for high probability of interaction insertion/deletion (*p* = 0.1) similar levels of connectance were achieved without losing species (Suppl. Fig. 3b), pointing to the role of the establishment of new interactions in driving increases in connectivity. Interestingly, trajectories stabilised for similar levels of connectance across different values of the parameter *p*, but the number of species varied. These results suggest that a central mechanism to sustain biodiversity possibly relies on building a core of well-connected species. This is also confirmed by the fact that the number of connected components in the network (the number of groups of nodes that did not share links with nodes outside of it) consistently converged to a single group (Suppl. Fig. 4).

Differences between trajectories under selection and neutral seems to be common across most topological properties analysed: in the absence of environmental perturbations there are significant differences while, if there are environmental perturbations, these differences are maintained only when interspecific competition is low. To provide further evidence, all networks under selection, except the one at high competition and high environmental perturbations, evolved their connectances towards a more skewed distribution of the degrees (*σ*(*degree*) Fig. 4, right column), being more disassortative for selection when environmental perturbations were absent and having similar values when perturbations were present (Suppl. Fig. 4). Nestedness is expected to increase for more disassortative networks and, indeed, when it was estimated with the measure of ecological overlap (namely, the mean fraction of shared mutualistic partners across species, see Methods [9]) it strongly increased along evolutionary time (Suppl. Fig. 4). However, when it was controlled by the expected nestedness of random networks with the same size and connectance using the NODFc measure (see Methods; [31]), it decreased for simulations under selection unless competition was high and environmental perturbations present (Fig. 4, middle column). Therefore, the disruptive nature of the evolutionary process we proposed limits our capacity to elucidate the role of nestedness on the final outcome of network assembly, a question discussed in more detail below.

**Figure 4:**
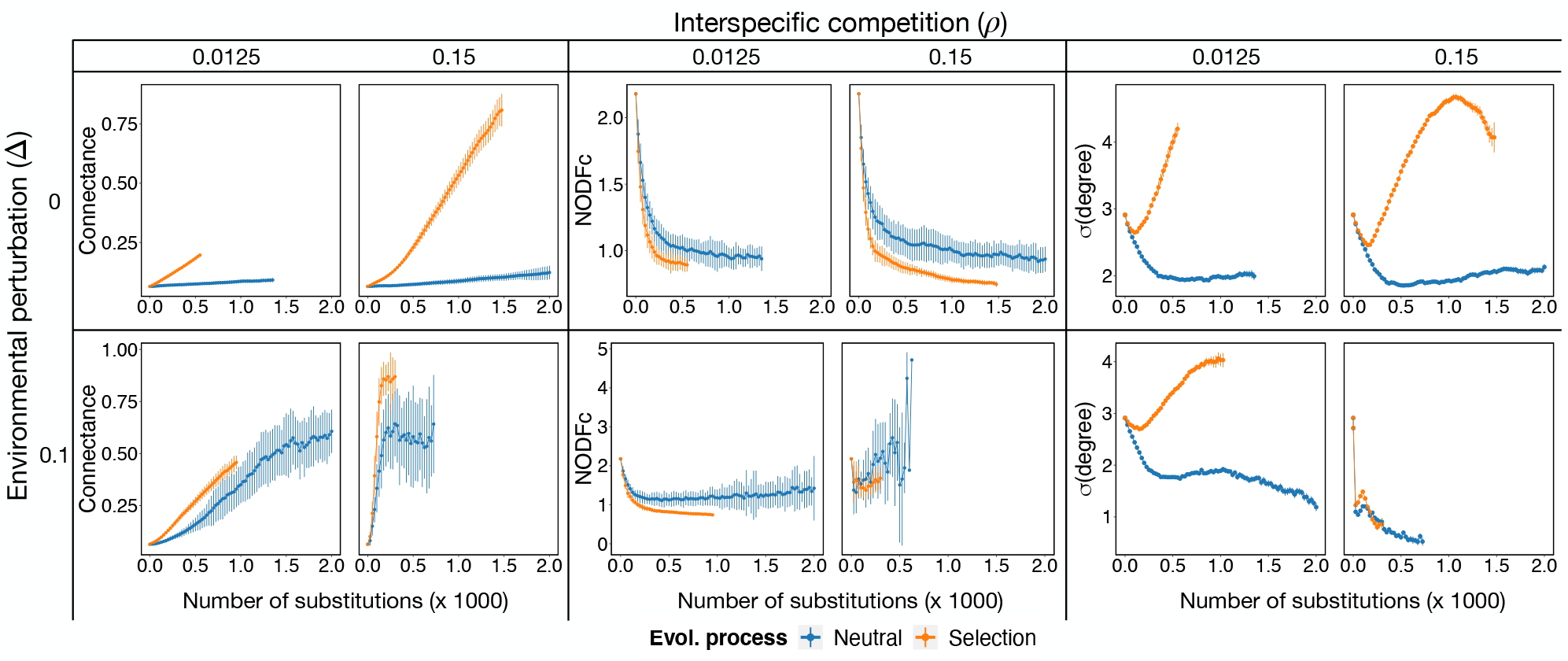
Evolution results in more connected networks that are less nested. Mean connectance (left), normalised nestedness (quantified as NODFc, middle) and standard deviation of species degrees (right) throughout eco-evolutionary trajectories (halted after stability in number of species is achieved over 350 evolutionary steps) for different values of environmental perturbation Δ (rows) and interspecific competition *ρ* (columns). Mean values (points) and standard deviation (vertical lines) are shown every 25 evolutionary steps (i.e. fixed mutation or substitutions) across 50 independent simulations for selection (orange) or neutral (blue) scenarios.

In summary, the disruptive evolutionary processes we built led to networks with fewer species, higher and unevenly distributed biomass, and higher, unevenly distributed connectances in the presence of selection, with nestedness having a secondary role in these networks.

### Evolution increases critical competition, favouring positive mutualistic effects

As shown above, the sign of mutualistic effects depends on the relative value of *ρ*_eff_ vs *ρ*_c_. Mutualistic interactions are detrimental for stability if *ρ*_eff_ > *ρ*_c_ and beneficial if *ρ*_eff_ < *ρ*_c_. Therefore, we considered the normalised distance of *ρ*_eff_ to *ρ*_c_, *D* = (*ρ*_c_ − ρ_eff_)/*ρ*_c_, defined between -1 and 1, to monitor the mutualistic regime, with *D* > 0 favouring and *D* < 0 being detrimental to stability (see Methods). Since evolutionary changes modify both *ρ*_eff_ and *ρ*_c_, we are interested in the mutualistic regimes that are favoured throughout the eco-evolutionary process we considered.

We observed that, independently of the environmental perturbations, the critical competition of the system tended to increase (Fig. 5, middle column). However, while for no environmental perturbations this increase was a consequence of the selected topological properties (with no overall increase for neutral simulations), in the presence of environmental perturbations it seemed to be a byproduct of the loss of species. For the effective competition, selection generated an overall reduction which, for neutral simulations, depended on the environmental perturbations. For both critical and effective competition we observed a shift within the first ∼ 500 substitutions aligned to changes in the topology of the mutualistic network, in particular to the variance of the degrees (compare Fig. 5 with Fig. 3).

**Figure 5:**
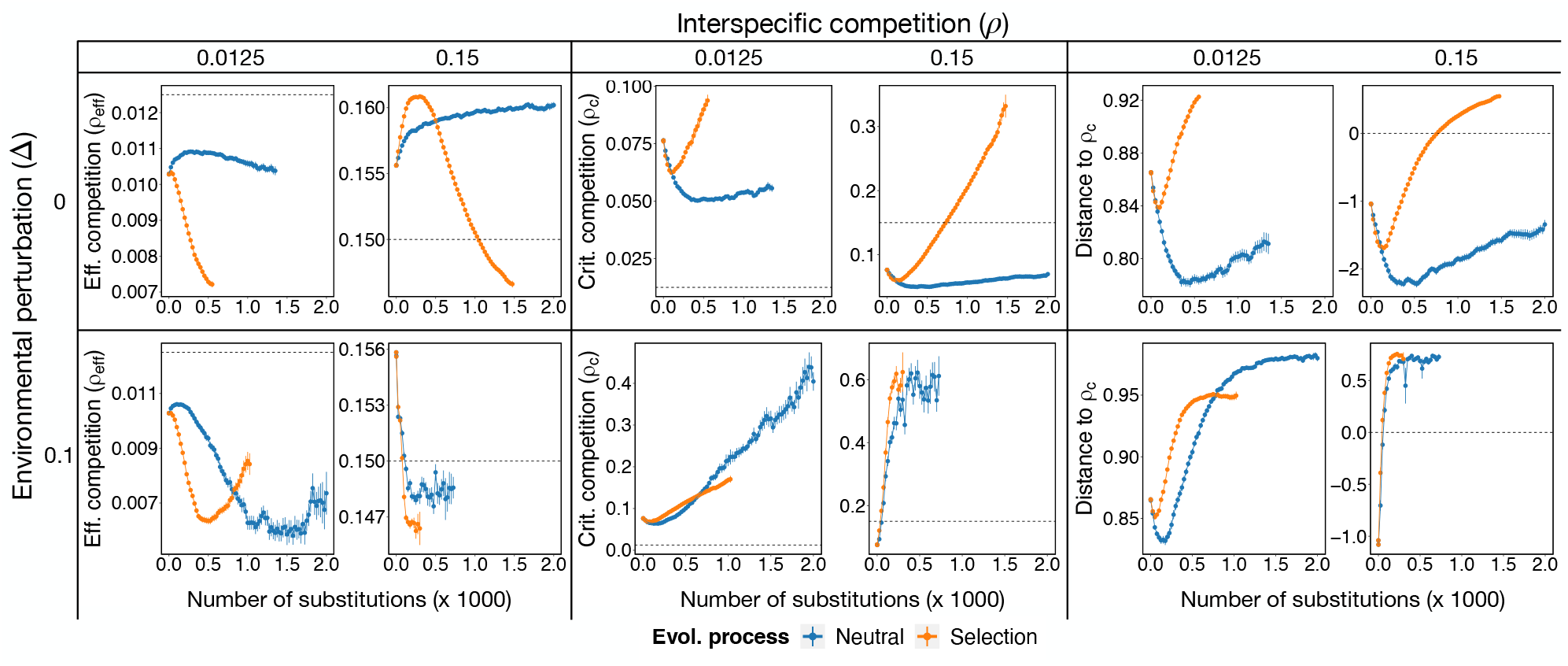
Evolution of effective and critical competition. Mean effective competition *ρ*_eff_, critical competition *ρ*_c_ (middle) and their relative distance *D* (right) throughout eco-evolutionary trajectories (halted after stability in number of species is achieved over 350 evolutionary steps) for different values of environmental perturbation Δ (rows) and interspecific competition *ρ* (columns). As a reference, *ρ* is also displayed with dotted lines in left and middle columns, and *D* = 0 in the right column. Mean values (points) and standard deviation (vertical lines) are shown every 25 evolutionary steps (i.e. fixed mutation or substitutions) across 50 independent simulations for selection (orange) or neutral (blue) scenarios. Evolution under selection always leads to systems with *ρ*_eff_ < *ρ* and *ρ*_c_ > *ρ*, implying *D* > 0 (*ρ*_eff_ < *ρ*_c_).

The complex interplay between both critical and effective competition explain differences in species loss between different parameter and selection regimes. In particular, for high competition we observed that the relative distance between effective and critical competition (*D*) run from negative to positive values (Fig. 5). When it was negative, mutualism had destabilizing effects and the strong increase in biomass observed for simulations under selection led to a dramatic loss of species. The system became stabilised, with no more species lost, when *D* became positive (crossing the dotted line in Fig. 5).

### Evolution increases structural stability, favouring biodiversity maintenance

We showed that the structural stability of the system at the beginning of the simulations sets the limits in which species loss will lie in an evolutionary process in the presence of environmental fluctuations. The structural stability, however, depends on the number of species and topological properties of the system, which vary throughout evolutionary trajectories. We were therefore interested in understanding how the structural stability itself evolved.

Notably, the structural stability of the system, estimated by the predicted critical perturbation, Δ_c_, experienced a long-term increase in the presence of selection (Fig. 6, left). While in some cases (e.g. neutral evolution, high competition and high environmental perturbations) this was a trivial consequence of having few highly connected species, this was not the case in other simulations (e.g. under selection, low competition, high environmental perturbations) where the number of species was above 50 (Fig. 3) and the connectance lower than 0.5, with a 3-fold increase in biomass (Fig. 6, left). Importantly, in several cases, the long-term increase occured only after a nuanced decrease, consistent with a similar shift observed for the effective competition, and suggesting that a topological rearrangement of the system that weakens its structural stability is a requisite to later enabling its increase.

**Figure 6:**
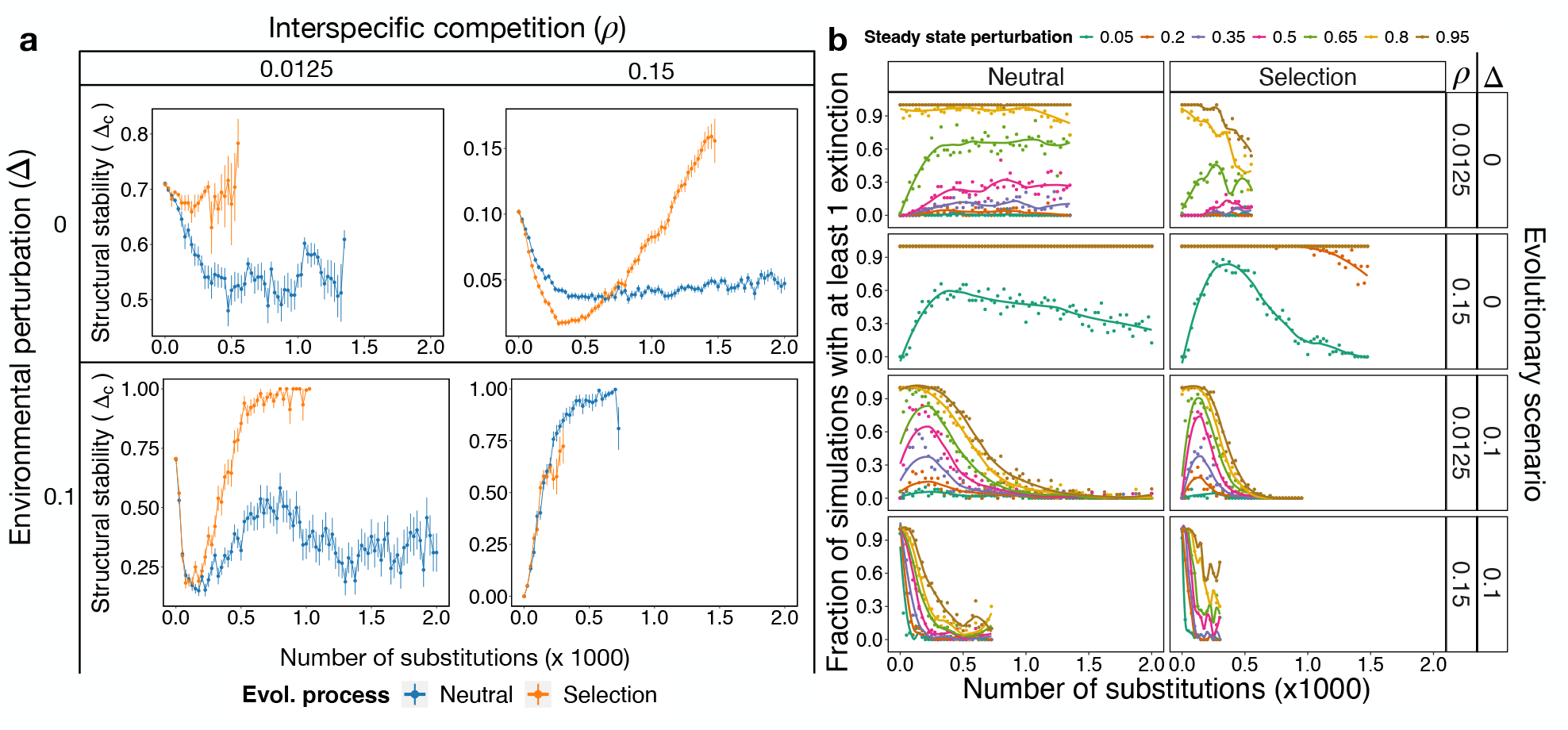
Selection increases structural stability across evolutionary time. (Left) Predicted structural stability, Δ_c_, for two interspecific competition values and without environmental fluctuations (upper row) and with fluctuations of amplitude Δ = 0.1 (lower row). (Right) Test of structural stability with simulations. Systems at steady state throughout the evolutionary simulations were tested numerically for structural stability every 25 evolutionary steps (x-axis) in the evolutionary scenarios of *ρ* and Δ combinations indicated in the right hand side. Each system was subjected to perturbations of amplitude Δ (shown on the top) and their dynamics investigated. Fifty replicates of ecological dynamics, corresponding to the 50 evolutionary replicates at the corresponding evolutionary time-points (i.e. number of substitutions), were simulated for each system and amplitude, and the fraction of simulations with at least one extinction is represented in the y-axis.

To validate structural stability predictions, we considered networks evenly distributed throughout the trajectories and tested their structural stability by perturbing the steady state found at the corresponding evolutionary time-point (see Methods). We considered different amplitudes of the perturbations and calculated the fraction of simulations in which at least one species went extinct.

Simulations were in agreement with predictions across all combinations of selection regime, competition and environmental perturbations (Fig. 6) with increasing Δ_c_ associated with a decrease in the fraction of simulations reporting extinctions. Importantly, we again observed a shift in which the system was less stable (increasing fractions of simulations with observed extinctions) to then become more structurally stable, especially for scenarios of high environmental perturbation and low competition (Fig. 6b), where the predicted structural stability of the system takes the larger relative fall (Fig. 6a).

Our results showed that, in the face of disruptive eco-evolutionary changes under selection, mutualistic systems will tend to lose species and to rearrange their topology to increase the critical competition above the effective one. This ensures that mutualism operates in the competition regime in which it favours stability.

## Discussion

Ecological and evolutionary perspectives on the importance of mutualisms for the emergence and persistence of biodiversity have revealed different facets of mutualistic networks assembly that influence their stability (reviewed in [20, 19]). Studies focusing on the evolution of mutualistic interactions have shown that interaction rewiring and speciation with inheritance are key ingredients to the emergence of structure in mutualistic networks ([21], [23]). On the other hand, those focusing on the stability effects of mutualisms on system dynamics at ecological time-scales, excluding evolutionary or assembly events (sometimes called internal stability [10]) have highlighted the crucial role of mutualistic interactions for community stability [32, 9, 16, 33, 15, 34]. These two angles have been, to a great extent, so far disconnected, possibly due to a lack of theoretical framework unifying ecological dynamics, evolutionary changes and invasions.

Indeed, most theoretical studies either explore the formation of patterns without taking into account the dynamics (e.g. [23]), or excluding the effects of environmental perturbations (e.g. [21]). Here, we contributed to fill this gap by studying how community coexistence becomes affected when evolutionary events are introduced and the ecological dynamics are explicitly simulated in the presence of environmental perturbations.

Our results showed that the equilibrium diversity of the eco-evolutionary process can be predicted from the estimation of the critical perturbation that the starting system can withstand. We observed predictability increased under selection, a result reported in evolutionary research [35]. To understand this prediction we should keep in mind that the critical perturbation is the maximum amplitude of environmental perturbations the system can cope with until at least one extinction is observed. On the other hand, the equilibrium diversity in the eco-evolutionary process was obtained after introducing repeated perturbations well below the critical value. Therefore, our results suggest a connection between repeated small perturbations and single large ones. A minimal amplitude of environmental perturbations seems to be a prerequisite to our ability to accurately predict outcomes of biodiversity, possibly to its role in overriding the effects of evolutionary changes.

We built on a well-studied population dynamic model for which the relationship between mutualism and biodiversity depends on the effective competition that species experience after accounting for all indirect pathways. By simulating evolutionary events in different environmentally-changing scenarios, we aimed to unveil the mutualistic regime (positive vs. negative for sustaining biodiversity) in which we could expect mutualistic systems to operate. We believe our approach provides an understanding of the ecological baseline on which other processes such as invasions or adaptation would play out (see below).

Our results suggest that, specially in the presence of selection, disruptive evolutionary change can drive mutualistic systems towards a regime in which mutualism supports biodiversity. We observed an increase in the total biomass of the system unevenly distributed across species, an increase in network connectance and, for selection, a decrease in the evenness of mutualistic degrees. Some of these patterns were a direct consequence of the loss of species (e.g. connectance in the presence of high environmental perturbations) but, in some simulation regimes, they were selected for.

The systematic increase of a well-connected core of species observed in our simulations seems to be aligned with results suggesting that connectance is underestimated in field experiments due to the typically short observations time-scales [36], and with the importance of core species in nested patterns [37]. It seems, however, to contradict findings pointing to the importance of an increase in modularity [22]. In our view, the discrepancy stands from the type of perturbations considered. While Sheykhali et al. [22] considered targeted attacks on specific species, we considered perturbations on model parameters, reflecting stochastic variation of environmental conditions. Directly targeting species abundances may make sense under some circumstances (e.g. human activity such as in fisheries or the use of certain types of antibiotics). In other scenarios, however, such as environmental conditions influencing species performance and, in turn, their abundances, the perturbations used in our study represent a more natural approach to model these conditions. Another explanation may come from differences in the selection strength since an increase in network connectance was an observed evolutionary outcome when selection was strong, while modularity was observed when selection was weak [38].

The main consequence of the topological and biomass changes we observed was an increase in the critical (*ρ*_c_), and an overall decrease in the effective (*ρ*_eff_), competition. The relationship between both quantities determine the mutualistic regime, being stabilizing when *ρ*_eff_ < *ρ*_c_ (Fig. 1b) and destabilising when the opposite is true. Importantly, in previous work we observed that in real networks a negative relationship between the critical competition and the number of species exists (Fig. 1d). Considering our results and assuming that real networks also operate in a regime in which mutualism positively influence stability, would imply that the larger the network is, the smaller must the effective competition be. This is an important finding, given the lack of experimental data to determine specific parameter values in theoretical modelling, a question that so far remains unsolved, particularly for competitive interactions [39]. This highlights the need to design sound experimental settings to measure and quantify specific demographic, environmental, and ecological (e.g. interaction strengths) dimensions of ecological systems. This research avenue to be prioritised in future studies aimed at understanding the drivers of the emergence and maintenance of biodiversity.

Our results suggest that determining the critical competition of mutualistic networks would allow theoreticians to determine regions of the parameter space describing their different behaviours, which can then be confronted with experimental data. Linking system properties with environmental disturbances can have important consequences for the monitoring of ecosystems’ health and the design of management interventions [40, 41]. As an example, we noted that, given an estimation of the interspecific competition, we could predict the region of environmental disturbance values where selection would be expected to have the strongest effects (see Suppl. Fig. 2).

Despite their simplicity, our results suggest connections with more complicated mechanisms. For example, the reduction in the effective competition seems to be aligned with the idea of mutualism having stabilising effects within evolutionary time-scales [42]. This also allows us to explain how other mechanisms explain coexistence when they operate by modulating competition, such as mutualistic-mediated adaptation to new niches [43], or competition–colonization trade-offs [44]. Further, we also observed that, although systems tend to become more structurally stable than the original network in the long-term, they achieved this state after a nuanced decrease in the structural stability. Therefore, environmental perturbations and evolutionary changes drive the system towards more vulnerable configurations, were some species were purged. This result provides a quantitative example of a qualitative metaphor described in the panarchy cycle [45], where it is suggested the need of a stage of ‘destruction’ to withstand perturbations. We observe, however, differences in the expected role of the connectance (destabilizing vs. stabilizing), the discussion of which remains beyond the scope of this work.

Our model has some limitations. Firstly, increasing the number of mutualistic interactions (i.e. beneficial effects) has no cost to a species, which may disproportionally favour more connected networks. A simple way to explore the effect of this was to reduce the probability of introducing/removing a link. In this setting, we found that increasing the probability of swapping interactions (at the expense of creating new ones) also induced an increase in network connectivity but through a different route: a decline in biodiversity. In our simulations, the sometimes abrupt decrease in the number of species and the increase in the connectance, rapidly reduces the sparsity of the interactions matrix, and hence the size of the ensemble of possible networks with similar levels of connectivity, making unlikely finding significantly nested patterns. This may explain contrasting results with the work of Suweis et al [21]. In that study, the authors reported that, when the introduction of new links was forbidden, a nestedness pattern emerged, in contrast with our results. The first difference with their work comes from the fact that they did not considered interspecific competition and, hence, it was not possible for mutualism to have disruptive effects [16], limiting the observation of extinctions. A second difference comes from the absence of environmental perturbations. Importantly, the authors did not report changes in the connectance nor in the richness of species throughout the evolutionary process, limiting thus the comparison with our results. However, they noted in the Suppl. Materials that: *‘the reduction of the effective ecosystem diversity due to species extinction that occurs while reaching stationarity, necessarily increases the nestedness of the corresponding interaction matrix because, on average, even randomly assigned matrices of smaller size are more nested’*. Therefore, in those cases in which simulations may have led to extinctions, an increase in the ecological overlap *v* (or *NODF*) would go hand-in-hand with increases in connectance, as we observed here. The use of a normalized measure of nestedness (such as the one we adopted here: *NODFc*) would have been required to determine if the observed increase in nestedness was significant in their simulations.

Secondly, we did not consider the introduction of new species [46, 25, 47]. The intended aim for this decision was to have a one-to-one comparison between evolutionary and structural stability. Since structural stability is defined as the maximum environmental perturbation a system can sustain before a species goes extinct, a direct comparison is to explore the evolutionary loss of species. We acknowledge that introducing new species may lead to other network topologies. For example, Cai et al. found similar results to ours with increased nestedness or modularity depending on model parameters [46]. Moreover, two mutualistic regimes were found in the comparison of stability between evolved and random networks when competition was varied, that could likely be explained by the critical competition threshold. Nevertheless, the pattern of variation between nestedness and modularity they observed when varying strength of mutualistic interactions may also be explained by the relative importance of nestedness and connectance in different parameter regimes [16]. Direct comparison with our results is not straightforward because they did not vary the connectance of the system. Combining their results with ours, we conjecture that each assembly event may reduce the distance between the effective and the critical competition, positioning the system at the edge of critical competition. Therefore, in our view, how a nested pattern emerges in an eco-evolutionary process with interspecific competition and environmental perturbations present is far from being understood. It may be needed processes such as adaptive foraging [48]], since it may favour specialist strategies, hence leading to more sparse and nested networks [47].

In summary, our work contributes towards an eco-evolutionary theory of ecological networks combining environmental fluctuations and evolutionary events, that consider mutualistic and competitive interactions together. Our results can have implications for conservation initiatives where the targets for conservation are not only species but their interactions. Especially since species interactions, and the corresponding effects on effective competition, can be strongly affected by different aspects of global change such as habitat loss and climate warming [49, 50, 51].

## Methods

### Ecological Community Dynamics

Following previous work [9, 16, 52], we model two groups of species: *S*^(P)^ plants and *S*^(A)^ animals. Species within each group compete with all others through fully connected competition matrices 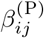 of Lotka-Volterra (LV) type. Species between groups interact via mutualistic interactions 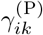, that are non-zero only if there is a link between *i* and *k* in the mutualistic network. Mutualistic benefits saturate at large abundances of the mutualistic partners providing the benefits [32]. The dynamics is hence governed by the following equations:

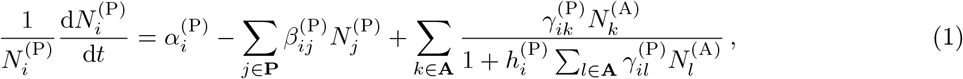

We only present the equations for plants, since those for animals can be exactly obtained by interchanging the superscripts P and A. In Eq. 1, *N*_*i*_ denotes the abundance the abundance of species *i, α*_*i*_ its intrinsic growth rate, and 1*/h*_*i*_ the maximum mutualistic growth rate of species *i*, which inverse, *h*_*i*_, can be interpreted as the handling time.

### Parameterization of the system

#### Competition and mutualistic matrices

In our model, we consider a competition matrix with different parameters for intra- (*β*_0_) and inter-specific competitions (*ρ*). The competitive matrix for plant species can be written as:

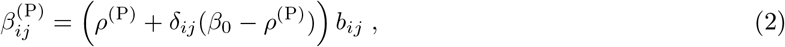

where *δ*_*ij*_ is Kronecker’s delta and *b*_*ij*_ are dimensionless parameter values uniformly distributed in [1 − ϵ_b_, 1 + *ϵ*_*b*_]. We parameterize mutualistic interactions as

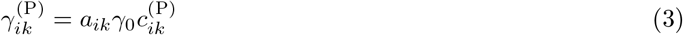

where *γ*_0_ measure the strength of mutualism with respect to competition, *a*_*ik*_ is the adjacency matrix of the mutualistic network, and *c*_*ik*_ are dimensionless parameter values uniformly distributed between 1 − ϵ_c_ and 1 + *ϵ*_*c*_ if *a*_*ik*_ = 1 and are zero if *a*_*ik*_ = 0.

#### Empirical network and metaparameters values

In our analyses, we consider as starting network a plant-pollinator mutualistic network with *S*^(A)^ = 46 animal and *S*^(P)^ = 47 plant species describing a rainforest mutualistic community comprising plants and their animal pollinators (insects and birds) from Cainama, Venezuela [53]. This network features approximately the same number of animal and plant species and connectance *κ* = 0.069 close to the median of a large set (*κ* = 0.09) of empirical networks available from the Web of Life database (https://www.web-of-life.es/).

The dynamical equations for all species in the system depend on seven meta-parameters: the direct interspecific competition parameters between plant (animal) *ρ*^(P)^ (*ρ*^(A)^) species ((0, 1)), the mutualistic handling times for plants *h*^(P)^(animals, *h*^(A)^), the mutualistic strength *γ*_0_, and the equilibrium abundances *N*^∗^. We used parameter values within the regime that allow us to simulate a weak-mutualistic facultative system, the regime in which topological trade-offs were clearly found in previous work [16]. In particular, we set *γ*_0_ = 0.15, *h* = 0.1 and *N*^∗^ = 1 for all species in the system. For each simulation, we parameterize the initial system randomly drawing a value for each parameter following a uniform distribution around its metaparameter value *X*: [*X − ϵ, X* + *ϵ*] with *ϵ* = 0.1*X*, except for *ρ* which remains constant, since we are interested in understanding the behaviour of the system across a wide range of interspecific competition values (see below). For simplicity, we set *ϵ*_*c*_ = *ϵ*_*b*_ = *ϵ*. Finally, the growth rates (*α*_*i*_) are determined once all the other parameters are drawn by solving the system at equilibrium, ensuring that they are positive and hence that the fixed point is feasible.

### Eco-evolutionary simulations

#### Evolutionary events and selection

The community dynamics model described above was coupled to an evolutionary model in which ecological interactions were subject to random change, extending the procedure described by Suweis et al. [21]. In this model, evolution proceeds in a series of timesteps at each of which a species *i*, on which random changes will be performed, is randomly selected from the whole set of species (including plant and animal species). Three possible mutation (or change) events might affect its interactions with other species in the system. With probability *p*, one of its mutualistic interactions (randomly selected) is swapped, and with probability 1 *− p* an interaction is either added (with probability *q*) or removed (1 *− q*) from its pool of existing interactions and randomly chosen. For simplicity, we chose *p* = *q* = 0.5. In the Suppl. Materials (Suppl. Fig. 3) we present results for simulations with *p* = 0.1 and *p* = 0.9 keeping *q* = 0.5. The strength of new interactions 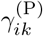 were drawn in the same way as the initial mutualistic interaction coefficients 3. Interspecific competitive interactions were not affected in the evolutionary process.

To model natural selection operating at the level of the species in a situation of sympatric speciation, evolutionary changes were accepted only if they were beneficial for the modified species, in which case we said that a substitution occured in the population. We defined this benefit as an increase in the population biomass at equilibirum of the modified species after simulating the ecological dynamics once the change was incorporated. Thus, changes were only accepted if the outcome of the ecological dynamics after change yielded a higher biomass at equilibirum for the mutated species compared to its biomass before the change, otherwise the change is rejected and the species goes back to its former interactions configuration. To ensure a fair comparison between evolutionary trajectories with and without selection, we considered trajectories with the same number of substitutions (i.e. rejected mutations are excluded).

#### Eco-evolutionary process

To simulate eco-evolutionary dynamics, we performed a two-step process in which ecological dynamics play out according to the dynamical equations (Eq.1) until the ecosystem reached equilibrium. The evolutionary steps described above were performed over the equilibrium state of the system resulting from the ecological dynamics.

To explore the effect of the strength of competition on the evolutionary trajectories of our networks, we performed simulations across a range of values of interspecific competition: *ρ* = {0.0125, 0.025, 0.05, 0.075, 0.1, 0.125, 0.15}. Intra-specific competition *β*_0_ was kept the same and equal to 1 throughout the duration of each single simulation. Further, to investigate the interplay between network evolution and the level of environmental perturbation, we explored the effect of varying the environmental perturbation on the system (i.e. Δ) by setting it to different values of Δ = 0, 0.025, 0.05, 0.1, where 0.1 corresponds to the lowest Δ_c_ value predicted for the network used in our simulations across all values of *ρ* used (i.e. *ρ* = 0.15, see Fig. 2b). We ran 50 simulations for each combination of environmental perturbation (Δ) and interspecific competition (*ρ*) values for both neutral and selection scenarios. During each simulation, at each evolutionary timestep (i.e. after ecological dynamics were integrated), species whose abundance fell below 10^−8^ were considered extinct and removed from the system. We considered that evolutionary stability was achieved when the number of species was maintained across at least 350 evolutionary timesteps. This procedure yielded a total of 2,800 evolutionary trajectories (7 *ρ*’s x ∼ 4 Δ’s x 2 selection regimes x 50 replicates) overall comprising ∼ 4 *×* 10^6^ population dynamics simulations (plus those performed for additional experiments considering different values of p, different initial networks, and structural stability analyses).

Numerical simulations of ecological dynamics were performed in Fortran, by integrating the equations (Eq. 1) using the Bulirsch-Stoer algorithm with adaptive step until convergence. This code was wrapped in R [54] to perform the evolutionary dynamics. Analyses of the simulation outputs were performed in R and plots generated with ggplot2 [55].

### Network properties

To assess changes in the state of the system along evolutionary trajectories we quantified total community biomass as the sum of the biomass of all species present in the commuity alongside a suite of network properties on the mutulistic interaction network. Standard network metrics such as the number of species *S* and connectance *κ* = *L/*(*S*^(P)^*S*^(A)^) (with *L* the total number of interactions in the network and *S*^(P)^and *S*^(A)^the number of species of plants and animals respectively) were complemented with additional structural properties.

The effective diversity (sometimes termed true diversity) was quantified as the exponential of the Shannon diversity. This is a convenient transformation since, if the relative abundances of species follows a uniform distribution, the effective diversity equals the *α* − diversity. The effective diversity will be lower otherwise, with more skewed distributions leading to lower values.

Nestedness, the extent to which the set of partners of specialist species are proper subsets of the partners of more generalist ones, was quantified using two metrics. Firstly, we defined it considering species ecological overlap *v*, which is the mean fraction of mutualistic partners [9]. For plant species:

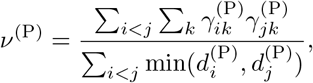

where 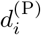 is the number of mutualistic links of plant *i*. A symmetric quantity is defined for animals simply exchanging the superscripts in the formula, with the total nestedness calculated as the arithmetic mean of both plants and animals quantities. Second, we considered the *NODF* metric proposed by Almeida-Neto et al. [56], which is qualitatively equivalent to *v* [21]:

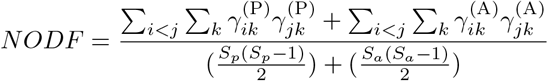

To enable comparison amongst networks of different sizes and number of interactions, we used a normalised version of *NODF* proposed by Song et al. [57]:

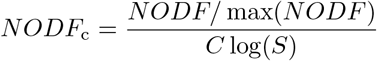

where max(*NODF*) is the maximum *NODF* nestedness value achievable in a network with the same number of species and links, and no disconnected species as the analysed network. *C* is the network connectance and *S* the geometric mean of the number of species in each level of the network. We used the function NODFc in the R package maxnodf [31] to calculate this measure.

Heterogeneity in the number of interactions across species was quantified as the standard deviation of the degree of the species in the network (*σ*(*degree*)). The degree of a species in the network is simply the number of mutualistic links it possesses.

Assortativity (*A*) is a network property that assess the tendency for nodes (i.e. species) to be connected to others with similar degrees [58].

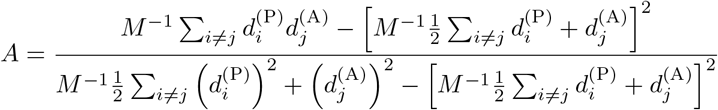

Lastly, the number of connected components in the network provides a measure of the extent to which the network is partitioned into isolated components not connected to one another. We counted the number of connected components as the number of groups of nodes that did not share links with nodes outside of it. If all nodes in the network are reachable from any other node, the number of connected components is 1. We used the R package igraph [59] to identify and count the number of connected components.

### Quantification of structural stability

#### Effective and critical competition

The linearization of the system governed by the system of equations 1 at equilibrium leads to the definition of the effective competition matrix, *C* (see Suppl. Materials and [9, 29] for a more general motivation of the approach). This matrix (*C*) encapsulates both the competitive interactions between species in the same group, either P or A, and their mutualistic interaction with species in the other group (see Suppl. Materials). The diagonalization of this matrix allow us to estimate the effective interspecific competition parameter for plant species (symmetric expressions are derived for animal species) as:

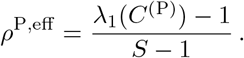

This parameter represents the mean-field competition experienced by a plant species in the system. Comparing *ρ*^P,eff^ to the “bare” interspecific competition parameter *ρ*, allows to assess the effect of mutualistic interactions on species. For example, if *ρ*^P,eff^ < *ρ* mutualism effectively reduces interspecific competition. In subsequent work [16], it was shown that mutualistic interactions reduce the interspecific competition of the system only if *ρ* is smaller than a threshold, termed critical competition, *ρ*^P,c^:

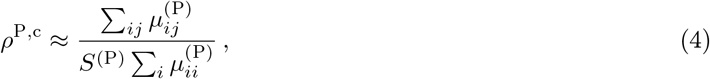

where 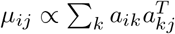, i.e. it is proportional to the mutualistic overlap between species in the same group. A full analytical derivation can be found in [16]. In Fig. 1 we recovered the *ρ*^P,c^ values obtained in [16] for a set of empirical networks retrieved from the web of life (www.web-of-life.es). The relationship between both quantities can be compactly expressed through the relative distance between the effective competition and the critical competition:

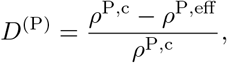

which is positive in the regime in which mutualism favours biodiversity, and negative otherwise.

#### Analytical prediction of structural stability

We follow the definition of structural stability proposed in Ref. [16], which considers the structural stability of a *specific fixed point* of a system. This differs from other definitions (e.g. [17]) that estimate the structural stability as the volume of the parameter space leading to feasible systems compatible with a given matrix of interactions, i.e. an estimation of *the volume all possible fixed points* [60]. Therefore, the definition we follow requires considering not only the interaction matrix but also the growth rates of all species and their realised abundances.

We model the effect of environmental perturbations of amplitude Δ on species growth rates as *α*_*i*_(Δ) = α_i_(Δ = 0)(1+Δr_i_), with *r*_*i*_ ∈ (− 1,1) being a random number drawn from a uniform distribution. We are interested in predicting the critical perturbation Δ_c_ in which at least one extinction was observed. This is our estimate of the structural stability of the system. In previous work [16] we showed that a good approximation of Δ_c_ can be obtained with the expression:

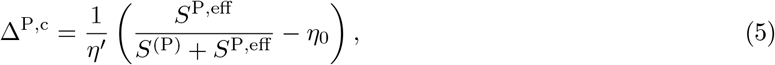

with symmetric equations for animal species. Hence, the system’s critical perturbation is the minimum found across both pools of species 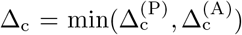. The term *S*_eff_ = (1-*ρ*_eff_)/*ρ*_eff_ represents a natural scale of biodiversity set by the effective competition *ρ*_eff_ and the quantities *η*^*′*^ and *η*_0_, which quantify the propagation of perturbations and the vulnerability of the system at the unperturbed state, respectively (see Suppl. Materials). Full details of the derivation of this expression is provided in Ref. [16]

#### Numerical quantification of structural stability

We verified that the analytical prediction of structural stability was consistent with the persistence observed in simulations. To this end, we considered systems at steady states every 25 substitutions throughout the eco-evolutionary trajectories. For each system at a steady state, we performed a random perturbation on the growth rates of amplitude Δ, and integrated the population dynamics of the system with the perturbed growth rates until a new steady state is reached, recording the number of observed extinctions. We repeated the same procedure for 50 random realizations of noise, and reported the fraction of simulations in which at least one extiction was observed. We investigated how this fraction changed for different values of Δ ∈ (0, 1).

## Acknowledgements

The authors thank Ugo Bastolla for valuable discussions. ML was supported by the French ANR through LabEx TULIP (ANR-10-LABX-41; ANR-11-IDEX-002-02), by a Region Midi-Pyrénées Project (CNRS 121090), and by the FRAGCLIM Consolidator Grant, funded by the European Research Council (ERC) under the European Union’s Horizon 2020 research and innovation program (grant agreement number 726176). APG was funded by a Ramón y Cajal Fellowship from the Spanish Ministry of Science and Innovation (RyC2021-032424-I), by CSIC intramural project 20232AT031 and by grant PID2022-139900NA-I00 (AEI/10.13039/501100011033/ FEDER, UE).

## Software availability

Source code developed to perform eco-evolutionary simulations and analyses are available at: https://github.com/computational-ecology-lab/eco-evolutionary-mutualistic-networks. A standalone Fortran code to run the population dynamics is available in URL: https://github.com/apascualgarcia/RatioDependent. Matlab code for the quantification of effective and critical competition and for predictions of structural stability is available in URL: https://github.com/apascualgarcia/structural_stability_prediction.

