## Supplementary Materials for "Structural stability determines evolutionary stability in mutualistic model ecosystems"

**3 Supplementary Figures** **8**

### 1 Supplementary Note: Evolutionary outcome in other regimes

#### 1.1 Evolutionary outcome for random networks

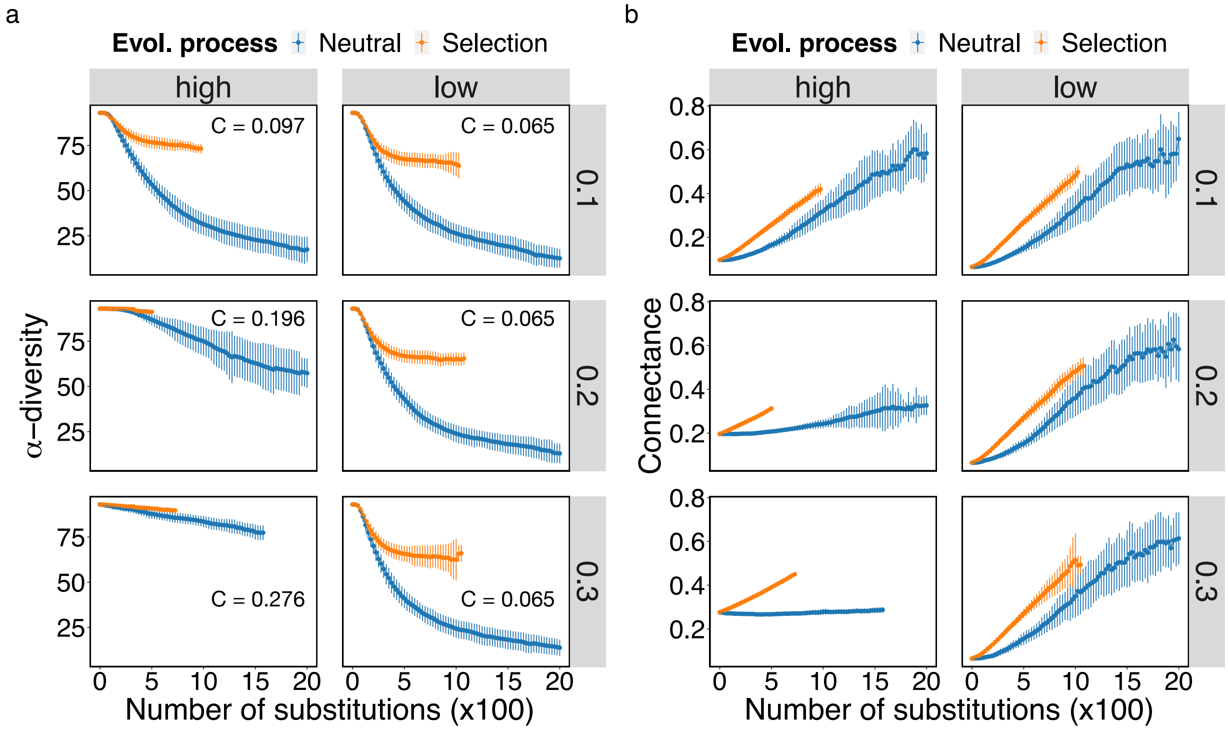

Figure 1: **Effects of initial network structure on evolutionary trajectories.** Mean (+/- standard deviation) $\alpha$-diversity (a) and connectance, b) throughout eco-evolutionary trajectories (halted after stability in number of species is achieved over 350 evolutionary steps) for initial networks with the same number of species as those explored in the main text (47 plants and 46 pollinators) but different structural properties. Nestedness ($\nu$) of the initial network was varied across three values $\{0.1, 0.2, 0.3\}$ (rows), while two values of connectance were selected for each value of nestedness: a high and a low relative value (columns). Connectance values are shown inside the corresponding plots in panel a. Note that the same values of connectance cannot be used because the lower the nestedness is, the lower the maximum value the connectance can attain. Mean values (points) and standard deviation (vertical lines) are shown every 25 evolutionary steps (i.e. fixed mutation or substitutions) across 50 independent simulations for selection (orange) or neutral (blue) scenarios. For these simulations the values of interspecific competition $\rho$ and environmental perturbation $\Delta$ were kept constant across scenarios and equal to 0.0125 and 0.1, respectively.

#### 1.2 Perturbations in the scale of the system's critical perturbation

In Main Text we convincingly showed that, for fixed environmental perturbations, systems with lower interspecific competition have higher evolutionary stability, as predicted by their structural stability. Nevertheless, we do not have experimental information about the interspecific competition realized in nature. Therefore, we asked what is the behaviour of evolutionary trajectories under environmental perturbations of different amplitude, where amplitudes were adapted to the scale imposed by the interspecific competition of each system. This allowed us to explore a wider range of environmental perturbations, and to understand how sensible systems with different interspecific competition are to relative changes in the environmental perturbations (instead of absolute values presented in Main Text).

More precisely, for a system with interspecific competition $\rho$, since we know that its critical environmental amplitude is $\Delta_c$, we simulated different evolutionary trajectories for perturbations $\Delta = \delta\Delta_c$ where $\delta \in (0, 1)$ is a scaling factor. We considered systems with $\rho = \{0.0125, 0.05, 0.1, 0.15\}$ and evolutionary trajectories with scaling factors $\delta = \{0,0.0125,0.05,0.1,0.25,0.5\}$.

We observed that selection led to systems with higher diversity except if $\delta$ =0, i.e. when no environmental perturbations were considered in the evolutionary process, where the neutral process preserves more diversity for higher competition (see Fig. 2a). In the presence of environmental perturbations, selection always preserved more diversity. Importantly, although we rescaled the environmental perturbations to make them equivalent throughout systems with different $\rho$, the final diversities were not the same. We observed that systems with higher $\rho$ were more sensitive, with their final diversity being smaller (Fig. 2a).

To connect with results presented in Main Text, we represented the difference in biodiversity between the simulations under selection and neutral, against the actual environmental noise under which the system was posited, i.e. $\Delta = \delta\Delta_c$ (Suppl. Fig. 2b). We found that the curves contained those presented in Main Text, as expected (Suppl. Fig. 2c). Interestingly, for each value of interespecific competition, there was a well defined maximum that peaked at a different amplitude of the environmental perturbations $\tilde{\Delta}$. We found an exponential relationship $\tilde{\Delta} \sim \exp(-\rho)$ indicating that for low competition, a small reduction of $\rho$ allowed to cope with large increases in the environmental perturbations $\Delta$ whereas, for high competition, large reductions in $\rho$ are required to cope with small increases in the environmental perturbations. Interestingly, for a system with known $\rho$ the value of $\tilde{\Delta}$ indicates the value of environmental perturbations in which the effects of selection will be stronger, a fact that may be useful to design interventions in ecosystems.

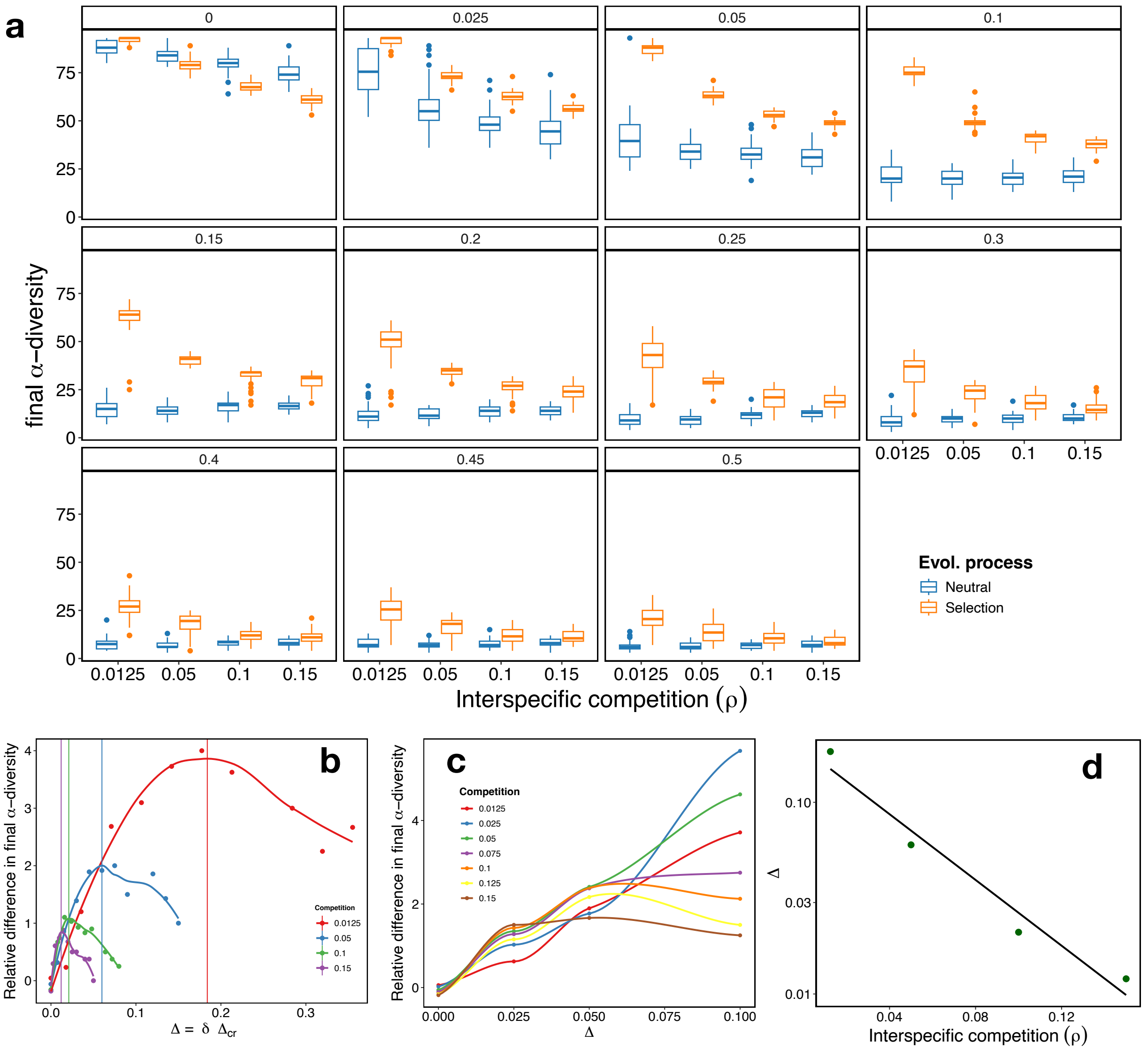

Figure 2: **Effects of the interplay between interspecific competition and the scaling of environmental perturbations on biodiversity.** a) Biodiversity of the eco-evolutionary process at equilibrium for different values of the interspecific competition $\rho$ and relative environmental perturbation $\delta$ (indicated in the title of the facets). Systems are under an actual environmental noise $\Delta = \delta\Delta_c$, were $\Delta_c$ are 0.71, 0.3, 0.16 and 0.1 correspond to $\rho$ values 0.0125, 0.05, 0.1 and 0.15, respectively. b) Relative difference in the final $\alpha$-diversity between simulations under selection and neutral, for different values of actual noise $\Delta = \delta\Delta_c$ and $\rho$. c) Same results as in b) in the region studied in Main Text. d) $\Delta$ value in which the maximum relative difference of biodiversity shown in b) was found, against the interspecific competition $\rho$. y-axis is in logarithmic scale, and a linear regression is displayed as a reference, indicating an exponential decay of $\Delta$ with increasing $\rho$.

#### 1.3 Effect of varying the probability of evolutionary events

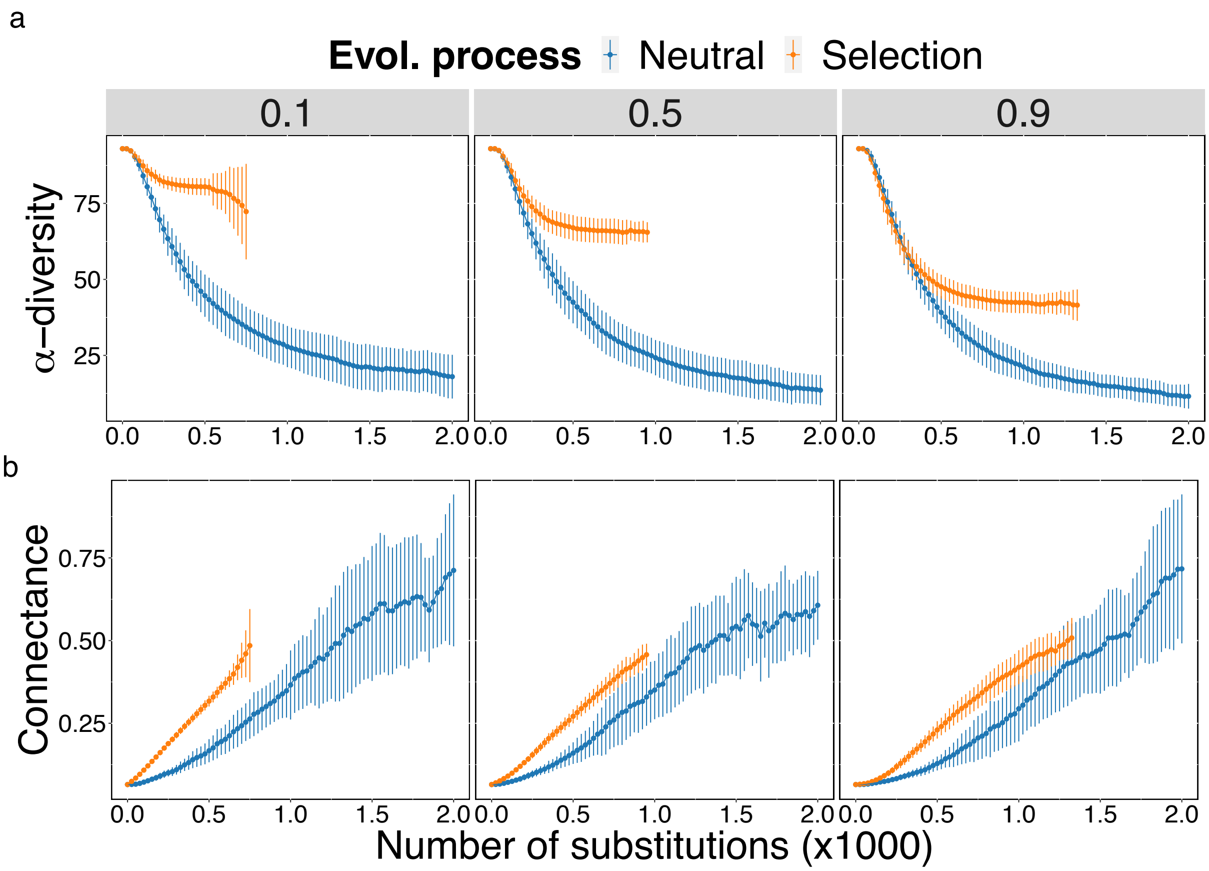

Figure 3: **Interaction swapping increases connectance at the expense of biodiversity in mutualistic networks.** Mean (+/- standard deviation) $\alpha$-diversity (a) and connectance b) throughout eco-evolutionary trajectories (halted after stability in number of species is achieved over 350 evolutionary steps) for different values of $p$ (probability of link swap during evolutionary events, columns). Mean values of each quantity (points) and standard deviation (vertical lines) are shown every 25 evolutionary steps (i.e. fixed mutation or substitutions) across 50 independent simulations for selection (orange) or neutral (blue) scenarios. In these simulations the environmental perturbation ($\Delta$) applied to the system was set to 0.1 whereas interspecific competition ($\rho$) was set to 0.0125.

### 2 Supplementary Note: Structural stability prediction and numerical quantification

#### 2.1 Equivalent Lotka-Volterra system and local stability

Close to a dynamical equilibrium, the dynamical stability for small perturbations of the abundances is determined by the equivalent Lotka-Volterra (LV) system

$$\frac{1}{N_i^{(P)}}\frac{\mathrm{d}N_i^{(P)}}{\mathrm{d}t} = \alpha_i^{\mathrm{eff}(P)} - \sum_{j\in\mathbf{P}} \beta_{ij}^{(P)} N_j^{(P)} + \sum_{k\in\mathbf{A}} \gamma_{ik}^{\mathrm{eff}(P)} N_k^{(A)} \,. \tag{1}$$

The effective interaction and growth rates parameters are obtained by differentiating the full dynamical equations of the full system (Eq. 1 in Main Text) at the equilibrium point, and they are

$$\gamma_{ik}^{\mathrm{eff}(P)} = \frac{\gamma_{ik}^{(P)}}{(1+z_i)^2} \tag{2}$$

$$\alpha_i^{\mathrm{eff}(P)} = \alpha_i^{(P)} + \frac{1}{h_i^{(P)}}\left(\frac{z_i}{1+z_i}\right)^2 . \tag{3}$$

$$z_i = h_i^{(P)} \sum_{l\in\mathbf{A}} \gamma_{il}^{(P)} \overline{N}_l^{(A)}$$

We see from this equation that for each species there are two regimes of parameters: weak mutualism, in which the equilibrium mutualistic benefit is far from saturation ($z_i \ll 1$) and strong mutualism, in which the saturation is reached ($z_i \gg 1$)

In previous work [1], we showed that the linearization of the system governed by Eqs. **??** can be described at steady-state, $\bar{N}$ by the equations:

$$\bar{N}^{(P)} = \left(C^{(P)}\right)^{-1} p^{(P)}; \qquad \bar{N}^{(A)} = \left(C^{(A)}\right)^{-1} p^{(A)} \tag{4}$$

where the effective competition matrices $C$ and the effective productivity vectors $p$ are defined as

$$C^{(A)} = \beta^{(A)} - \gamma^{\mathrm{eff}(A)}\left(\beta^{(P)}\right)^{-1}\gamma^{\mathrm{eff}(P)}; \qquad p^{(A)} = \alpha^{\mathrm{eff}(A)} + \gamma^{\mathrm{eff}(A)}\left(\beta^{(P)}\right)^{-1}\alpha^{\mathrm{eff}(P)} \tag{5}$$

and analogous for plants. The effective competition matrix represents the interactions between species in the same group, either P or A, both due to their direct interaction (in this case, competition) and to their interaction with species in the other group (in this case mutualism). The effective productivities, describe the influence of competitive and mutualistic interactions on the effective growth rates $\alpha^{\mathrm{eff}(A)}$.

#### 2.2 Structural stability prediction

We present here a simplified description of the derivation of the structural stability prediction presented in Main Text for completeness. The full derivation is presented in [2]. To simplify the notation, in the following we omit the superscripts for plants and animals since the equations are symmetric. In Main Text, whenever a quantity defined for each pool is presented, we select the one for plants by default.

Since we are interested in the structural stability of a given fixed point, we first estimate the vulnerability $\eta_i$ of each species at that particularly state, which can be calculated using the expression

$$\eta_i = 1 - \frac{p_i}{v_i^1 p^1}, \tag{6}$$

where $p_i$ is the effective productivity of species $i$, $v_i^1$ is the component $i$ of the main eigenvector of the effective competition matrix (see above), and $p^1$ is the projection of the productivity vector onto the main eigenvector, i.e. $p^1 = \sum_j p_j v_j^1$. Intuitively, this term estimates how vulnerable a species is by computing how far its productivity is from the main eigenvector of the effective competition matrix. This is justified by the fact that the more parallel the productivity vector and the main eigenvector of the effective competition matrix are, the more structurally stable the system is. In previous work it was shown that, for large $\Delta$, the most vulnerable species is always the same and, thus, to estimate the vulnerability of the system we can focus on the most vulnerable species, i.e. $\eta = \max_i(\eta_i)$.

Additionally, as shown in [3], a necessary condition for the feasibility of a system is that the vulnerability of all species falls under a threshold given by

$$\eta \leq \eta_{\rm c} = \frac{S_{\rm eff}}{S + S_{\rm eff}}. \tag{7}$$

This expression directly connects the feasibility of the system with both its biodiversity and with the effective competition present in the system, since the larger $S$ or $\rho_{\rm eff}$ are the most difficult is to fulfill the condition.

In the presence of an environmental fluctuation of amplitude $\Delta$, the growth rates $\alpha$ are affected and, in turn, the effective productivities $p_i = p_i(\Delta)$ and vulnerabilities $\eta = \max_i(\eta_i(p_i(\Delta)))$. For large amplitudes the perturbed vulnerabilities are approximately linear with $\Delta$,i.e. $\eta(\Delta) \approx \eta_0 + \eta'\Delta$, where $\eta_0$ is the vulnerability value for the most vulnerable species at the unperturbed state and $\eta'$ is the propagation of perturbations. The quantities $\eta_0$ and $\eta'$ are estimated for two sufficiently large perturbations around $\eta_{\rm c}$ as

$$\eta' = \frac{\eta(\Delta_1) - \eta(\eta_0)}{\Delta_1 - \Delta_0}, \qquad \eta_0 = \eta(\Delta_0) - \eta'\Delta_0$$

with $\Delta_0 = \eta_{\rm c} - 0.05$ and $\Delta_1 = \eta_{\rm c} + 0.05$. Once these quantities are obtained, the linear approximation of $\eta(\Delta)$ is substituted in Eq. 7 and solved for $\Delta$. The critical perturbation is obtained when the equality in Eq. 7 is fulfilled:

$$\Delta^{\rm P,c} = \frac{1}{\eta'}\left(\frac{S^{\rm eff(P)}}{S^{\rm (P)} + S^{\rm eff(P)}} - \eta_0\right), \tag{8}$$

Since the perturbations are unknown in advance, we followed a semiempirical strategy in which random perturbations of fixed amplitude $\Delta$ are drawn, generating a distribution of values for the right-hand side of Eq. 8. We then take as $\Delta_{\rm c}$ the amplitude which equals the median of that distribution.

### 3 Supplementary Figures

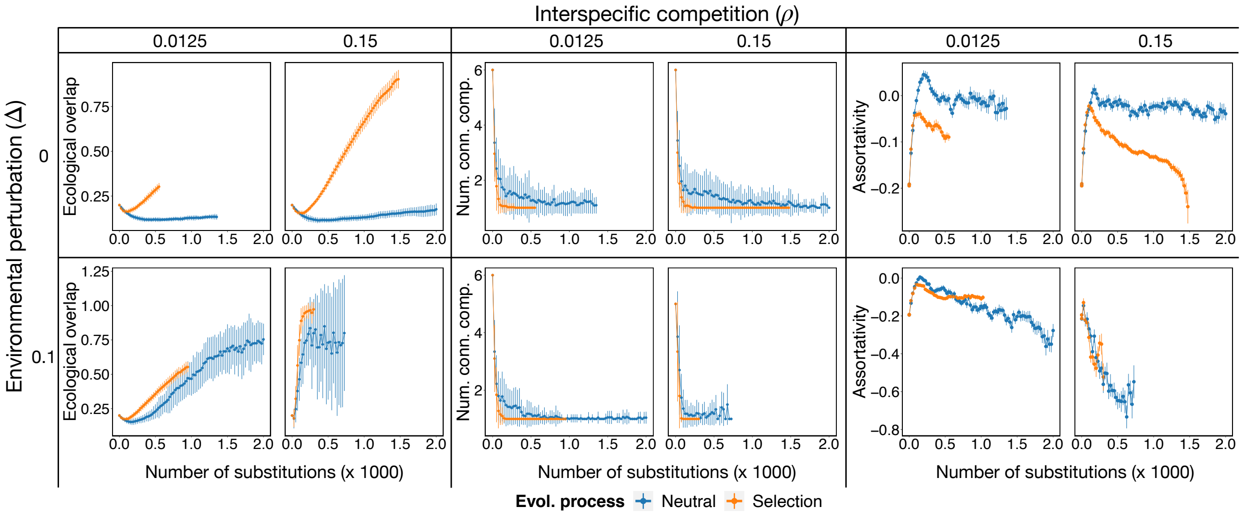

Figure 4: **Evolution of structural properties.** (Left) Nestedness (ecological overlap $\nu$), number of connected components (middle) and assortativity (right) throughout eco-evolutionary trajectories (halted after stability in number of species is achieved over 350 evolutionary steps) for different values of environmental perturbation $\Delta$ (rows) and interspecific competition $\rho$ (columns). Mean values (points) and standard deviation (vertical lines) are shown every 25 evolutionary steps (i.e. fixed mutation or substitutions) across 50 independent simulations for selection (orange) or neutral (blue) scenarios.
